# Establishing Preclinical Withdrawal Syndrome Symptomatology Following Heroin Self-Administration in Male and Female Rats

**DOI:** 10.1101/2020.03.02.974006

**Authors:** Cassandra D. Gipson, Kelly E. Dunn, Amanda Bull, Hanaa Ulangkaya, Aronee Hossain

**Affiliations:** Department of Family and Community Medicine, University of Kentucky, Lexington, KY; Behavioral Pharmacology Research Unit, Johns Hopkins University, Baltimore, MD; Department of Psychology, Arizona State University, Tempe, AZ

**Keywords:** Opioid, Withdrawal, Timecourse, Heroin, Self-Administration, Sex Differences

## Abstract

Opioid use disorder (OUD) is a significant health problem, and understanding mechanisms of various aspects of OUD including drug use and withdrawal is important. Preclinical models provide an ideal opportunity to evaluate mechanisms underlying opioid withdrawal. Current models are limited by their reliance upon forced opioid administration, focus on the acute (and not protracted) syndrome, and exclusion of females. In this study, male and female rats self-administered heroin (maintenance dose of 12.5 μg/kg/infusion) opioid withdrawal following abrupt discontinuation was measured. In Phase 1, acute withdrawal symptoms were rated in male and female rats at 0, 16, 48, and 72 hrs following the last self-administration session. Total somatic signs increased until 48 hrs (predominantly in females), and heroin intake positively correlated with total somatic signs at the 48 and 72 hr timepoints. Measures of hyperactivity and anxiety-like behavior increased by 16 and 48 hrs, respectively. In Phase 2, symptoms were assessed at baseline, acute, and protracted (168 and 312 hrs after self-administration) timepoints in a subset of male and female rats from Phase 1. The total number of somatic signs did not differ across timepoints, though females displayed significantly higher body temperature at all timepoints compared to males, indicating sex-specific protracted withdrawal symptomatology. These data provide a thorough characterization of rodent opioid withdrawal symptomatology following self-administration and abrupt discontinuation that serve as a foundation for future studies designed to mimic the human experience, and demonstrate the importance of characterizing acute and protracted withdrawal with sex-specificity in preclinical models of opioid self-administration.

## Introduction

Use and misuse of opioids is a major global health issue that results in high levels of mortality and morbidity, and significantly decreases life expectancy (Dowell et al., 2017). Opioid use disorder (OUD) is characterized by both a persistent desire to use the drug as well as a desire to suppress the aversive symptoms of opioid withdrawal. The specific incidence, timecourse, and severity of opioid withdrawal symptoms in humans varies depending on the opioid being used and the duration of use, and consists of both an acute syndrome that lasts approximately 7 days and a protracted syndrome that can last for several weeks (Jasinski, 1981). In humans, symptoms of acute withdrawal begin to emerge 4-7 hours (short-acting) or 24-36 (long-acting) after the final opioid exposure. Common acute symptoms include muscle aches, nausea, diarrhea, vomiting, hot and cold flashes, piloerection, lacrimation, and acute anxiety, and common protracted symptoms include persistent craving, sleep impairment/insomnia and mood disruption. For both syndromes, symptoms emerge at myriad times, reach peak severity at different rates, and last for variable durations. Thus far, efforts to reliably predict opioid withdrawal severity in humans have been largely unsuccessful, with studies attributing differences in withdrawal presentation to several different demographic and drug use characteristics.

A recent review outlined the numerous species that are used for preclinical evaluation of substance use disorders (Smith, 2020), including invertebrate, rodent (de Guglielmo et al., 2017), and nonhuman primate models (Banks et al., 2017; Katz, 1986; Schwienteck, Negus, & Banks, 2019). These models provide a method by which correlates and underlying mechanisms of different opioid withdrawal symptoms can be precisely evaluated, and have been used in the past to directly support clinical evaluation of various withdrawal pharmacotherapies. Rats provide an ideal platform for this research because their opioid self-administration behavioral paradigms are well-established and lead to significant rates of voluntary drug self-administration (Ahmed, Walker, & Koob, 2000; Bossert et al., 2015; Mavrikaki et al., 2017). In addition, nodes within the mesocorticolimbic reward circuitry are conserved between rats and primates (Heilbronner et al., 2016), lending translational value to rats as a model species. The vast majority of preclinical rodent studies induce opioid physical dependence through forced administration of an opioid (e.g., subcutaneous morphine pellet, injection, or infusions) and then precipitate withdrawal using the fast-acting opioid antagonist naloxone (e.g., see Dunn et al., 2019 for review). Common withdrawal symptoms assessed in these studies include body shakes, chewing, diarrhea, digging, escape attempts, eye blinks, foot licks, genital licks, grooming, head shakes, jumping, piloerection, ptosis, rearing, teeth chatters, writhing, and change in temperature.

Although these studies provide a valuable opportunity to assess species-specific neural mechanisms of opioid withdrawal, their methods do not reflect real-world human consumption patterns. For instance, engendering opioid physical dependence with subcutaneous morphine implants (Ary & Lomax, 1976; Cox, Ary, & Lomax, 1975) or morphine subcutaneous injections (el-Kadi & Sharif, 1998) is able to produce quantifiable opioid dependence but it is not clear whether the withdrawal produced from this exposure type will resemble the withdrawal syndrome produced from self-administration of opioids. Self-administration allows an animal to stop responding for drug once they experience negative effects, which corresponds to human drug self-administration. As a result, the level of dependence produced through forced administration could be far higher or lower than what might have occurred in a self-administration paradigm, thus calling into question the generalizability of that model to human subjects. In addition, using a bolus dose of an opioid antagonist to precipitate withdrawal produces a brief and high magnitude withdrawal syndrome but that severity of withdrawal may not reflect real-world conditions wherein withdrawal symptoms emerge spontaneously and in a more mild manner following opioid discontinuation (Pinelli & Trivulzio, 1997; Zhang et al., 2016).

A recent review of preclinical and human research on novel medications for opioid withdrawal management revealed the majority of drugs that showed a positive signal in animals did not translate into human research or failed in human trials (Dunn et al., 2019). These studies primarily relied upon the forced administration/precipitated withdrawal method, and it is possible this approach produced a high magnitude withdrawal that was sensitive to medical intervention but failed to generalize to the different course of withdrawal observed within humans. Moreover, since preclinical studies have traditionally precipitated withdrawal for an acute period (e.g., Papaleo & Contarino, 2006; Hodgson et al., 2010; Radke et al., 2013, Ramesh et al., 2013), they rarely examine withdrawal during the protracted period that is of major concern for human opioid withdrawal management. We know of no preclinical studies that have reported on the natural history of spontaneous opioid withdrawal in animals whose dependence is the result of their own heroin self-administration behavior. Further, despite growing recognition about the importance of examining potential differences in withdrawal presentation in men and women (Lewis, Hoffman, & Nixon, 2014), these domains have not yet been well-integrated into preclinical and human withdrawal examinations. This reflects a larger concern about the inequity of sex and gender research as it pertains to the opioid crisis; the Fast-Track Action Committee (FTAC) on (the) Health Science and Technology Response to the Opioid Crisis recently published concerns that women have been largely excluded from research on OUD mechanisms and treatment strategies (Becker & Mazure, 2019).

The need for refined models that facilitate good translation from animal to human samples has been recently outlined in a National Institute on Drug Abuse (NIDA) Request for Information (RFI) to “examine the translational value of current animal models” (also see Smith, 2020). Optimal preclinical models will mimic the human experience while providing an opportunity to rigorously examine opioid withdrawal mechanisms and interventions in highly-controlled and precise ways. Establishing the natural history of an ecologically representative model of spontaneous opioid withdrawal in animals will facilitate research on new potential pharmacotherapies and interventions for opioid withdrawal management. For these reasons, the present study aimed to characterize the opioid withdrawal syndrome timecourse and severity in male and female rats that were made physically dependent upon opioids following a standardized heroin self-administration paradigm and then underwent spontaneous opioid withdrawal following opioid discontinuation. Withdrawal ratings were also collected during an acute (Phase 1) and protracted (Phase 2) period. The goal of these data is to provide a foundation upon which methods for assessing opioid withdrawal in rodents can be standardized, to support improved generalization across preclinical studies and improve opportunities for translation between preclinical and human samples. We hypothesized that rats would show different patterns of expression throughout the acute and protracted heroin withdrawal periods (consistent with the human expression of withdrawal). We further hypothesized that the pattern of symptom emergence in both acute and protracted periods would be sex-specific, with withdrawal symptomatology being exacerbated in females compared to males. We also hypothesized females would show faster heroin self-administration acquisition and greater withdrawal symptomatology than males, based upon evidence from the human literature that women show more rapid transition from occasional to chronic opioid use than men (Becker & Koob, 2016; Greenfield et al., 2007), and show greater withdrawal response than men (Becker & Chartoff, 2019).

## Methods and Materials

### Animals

The number of animals per sex was determined by *a priori* power analyses. 19 male and 11 female Long Evans rats (200 – 250 g) were utilized in the current study. Of these animals, 7 were bred in-house as wildtypes of a transgenic rat breeding colony and 23 additional animals were purchased from Envigo. The number of animals purchased was determined by litter sizes in order to achieve appropriate power. The animals that were bred in-house were from 3 different litters, each with different male and female breeders and were comprised of ChAT::cre positive males (Long Evans-Tg(ChAT-Cre5)5.1 Deis; purchased from RRRC (RRRC #658; Columbia, MO)) and Long Evans wildtype females purchased from Envigo. All animals utilized in the current study were ChAT::cre negative males or females.

Rats were housed individually, handled daily, and were on a 12-hour reverse light cycle with *ad libitum* access to food and water prior to initiation of self-administration procedures. Phase 1 animals (19 males, 11 females) were assessed for evidence of acute withdrawal and then a subset of these animals (8 males, 8 females, from the Envigo lines) were assessed for protracted withdrawal symptoms during Phase 2 of the study. All animal use practices were approved by the Institutional Animal Care and Use Committee of Arizona State University (ASU) and wildtype animals from a breeding colony were used for this study consistent with the guiding principles for ethical use of animals in research (the three “R”s of research, including reduction, replacement, and refinement). No animals were omitted from the study, and all animals completed 14 self-administration sessions.

### Apparatus

Experimental sessions occurred in operant chambers (MED Associates, St. Albans, VT) within sound-attenuating chambers with ventilation fans. Two levers, one active and one inactive, were presented at each session. Above each lever was a stimulus light and each chamber contained a house light. Infusion pumps (MED Associates) were located outside of each chamber. Experimental events were recorded by MED-PC software (MED Associates) on a computer in the experimental room.

### Surgical Procedures

At post-natal day (PND) 60 and five to six days prior to undergoing food training, animals were implanted with a catheter in the jugular vein as described previously (Powell, et al., 2019). Briefly, animals were anesthetized with intramuscular (IM) ketamine (80-100 mg/kg) and xylazine (8 mg/kg, IM). A small incision was made in the chest to isolate the jugular vein, and a second small incision was made between the scapula. A 13 cm catheter (BTPU-040; Instech, Plymouth Meeting, PA) with an anchor made from laboratory tubing (Silastic®, Dow Corning, Midland, MI) 2 cm from one end was tunneled subcutaneously between the incisions, with the anchored side inserted into the jugular vein. The dorsal end of the catheter was attached to an indwelling back port (Instech, Plymouth Meeting, PA) using dental cement (SNAP; Parkell, NY or Ortho-Jet; Lang Dental, IL). Following successful placement of the catheter, post-operative cefazolin (100 mg/kg, intravenous (IV)) and meloxicam (1 mg/kg, subcutaneous) were administered for 7 and 3 days, respectively. Catheter patency was maintained throughout the study by infusion of 0.1 mL heparin saline (100 USP units/ml, IV) administered daily.

### Food Training

After implantation of jugular vein catheters, animals were food restricted to 20 g of chow for 5 hours and responding on the active lever was reinforced by the delivery of food pellets in the apparatus described above. Food training sessions lasted 15 hours and the houselight remained illuminated throughout training. Upon completion of a fixed ratio-1 (FR1) schedule of reinforcement on the active lever, a 45 mg food pellet (BioServ®, Flemington, NJ) was delivered into a food receptacle inside the operant chamber. No consequences were produced by pressing the inactive lever. Animals with an active to inactive lever press ratio ≥ 2 were removed from food restriction, and kept in their home cage until heroin self-administration began the following day. Any animals that did not reach this criterion underwent a second food training session the following day. All animals met the food training criterion after 1-3 sessions. If animals needed re-food training, they received 20g of chow following their first food training session, and then were given one day off to sit in their homecages and rest during their inactive phase. If needed on the following day, animals went into their second food training session at 5PM. Occasionally, chow was left over from the previous day’s feeding, and this would be taken away by 12PM on the second day. Following their second food training session, animals were again fed 20g and started self-administration the next day. Notably, 3 animals required a third food training session (this was the maximum number of food training sessions needed), but eventually all animals met criteria and moved into self-administration. Animals were food restricted throughout the duration of experimentation.

### Self-Administration Procedures

Heroin self-administration began one week after surgery with a fixed ratio (FR)-1 schedule of reinforcement. A step-down dose regimen was implemented to expedite acquisition of self-administration, as previously published (Fuchs & See, 2002; Shen et al., 2011). Infusions of 50 μg/kg/infusion heroin were delivered across 3 s in 0.05 mL following a response on the active lever for 2 sessions. The following 2 sessions consisted of a 25 μg/kg/infusion heroin dose. The maintenance dose of heroin, which was available from sessions 5-14, was 12.5 μg/kg/infusion. Lights above both levers were illuminated and a tone (2900 Hz) sounded simultaneously with drug infusion. Upon completion of the infusion, the lights and tone ceased, and the house light was illuminated for a 20-s timeout period, during which active lever responses were recorded but produced no consequences. An inactive lever was present at all times, but activation of the lever produced no consequences. Sessions lasted 3 hrs. Animals were required to complete 14 total sessions of heroin self-administration prior to measurements of withdrawal.

### Somatic Signs of Withdrawal

The following sixteen somatic signs of withdrawal were assessed: body shakes, chewing, diarrhea, digging, escape attempts, eye blinks, foot licks, genital licks, grooming, head shakes, jumping, piloerection, ptosis, rearing, teeth chatters, and writhing were measured at 0, 16, 48, and 72 hrs post heroin self-administration (Phase 1) and 0, 16, 48, 72, 168, and 312 hours post heroin self-administration (Phase 2; see Figure 2A for a timeline of procedures). As subset of animals were utilized for additional baseline and protracted timepoints (Phase 2). Baseline ratings of symptoms were collected three days prior to jugular catheter surgery. These specific withdrawal symptoms were selected for evaluation because they are commonly reported in preclinical examinations of opioid withdrawal and also correspond roughly to human symptomatology. Symptoms were quantified by placing rats in a clear Plexiglas chamber (9.6 x 9.6 x 14.6in; L x W x H) and recording behaviors for 10 minutes (following a 5 min habituation period). Experimenters scored withdrawal signs from the video recordings using a standardized scoring protocol.

### Physiological Signs of Withdrawal

In addition to somatic signs of withdrawal, point prevalence measurements of body temperature were collected via rectal thermometer prior to placing the animal in the chamber at each timepoint.

### Drugs and Chemicals

Diacetylmorphine (Heroin; Cayman Chemical, Ann Arbor, MI, USA) was dissolved in 0.9% saline. Ketamine (Akorn Animal Health, Lake Forest, IL), xylazine (Akorn Animal Health, Lake Forest, IL), cefazolin (Qilu Pharmaceutical, Shandong, China), meloxicam (Norbrook Inc., Overbrook, KS), and heparin (Sagent Pharmaceuticals, Schaumburg, IL) were administered as discussed above.

### Data Analysis

The primary goal of these studies was to characterize heroin self-administration and subsequent spontaneous withdrawal symptomatology among male and female rats. Analyses were identical across Phases 1 and 2 unless otherwise noted.

Heroin self-administration acquisition in all animals was assessed using a one-way repeated measures ANOVA with Bonferonni correction, comparing heroin infusions on each session to session 1. An omnibus three-way ANOVA treating lever and session as within- and sex as between-subjects factors was then conducted to evaluate acquisition rates within male and female samples. Rate of acquisition of heroin self-administration was then re-assessed within each sex subgroup using repeated measures two-way ANOVAs with lever and acquisition session as within-subjects factors. Rate of acquisition was then assessed within each sex group by comparing the number of infusions received across sessions using both repeated measures ANOVA and linear regression analyses (Gipson et al., 2011). The total number of heroin infusions was compared across sex groups using a one-tailed *t*-test and significant interactions were explored using Bonferroni-corrected post-hoc comparisons.

Spontaneous somatic signs of heroin withdrawal were analyzed by summing the number of expressions observed for each symptom at each timepoint. For ease of interpretation, withdrawal symptoms are presented in categories that represent the following general domains: anxiety/escape behaviors (digging, jumping, rearing, and escape attempts), body temperature (measured in °C, teeth chattering), gastrointestinal symptoms (diarrhea, defecation), hyperactivity (chewing, grooming, foot licking, genital licking), pain/hyperalgesia/ ptosis (writhing), and tremors/shaking (body, head shakes). These categories are based on meaningful clusters that have been identified within the human clinical literature (Dunn et al., 2019). Withdrawal severity (operationalized as number of presentations within each 10-minute scoring period) and total score (operationalized as total number of symptoms with >1 incidence during each 10-minute scoring period), as well as the physiological rating of body temperature were evaluated using separate repeated measures one-way ANOVA, with time as a within-subject repeated measure. A mixed model ANOVA was also conducted that treated sex as a between-subjects factor and timepoint as a within-subjects factor, and Bonferroni-corrected post-hoc comparisons were conducted when appropriate. To assess the degree to which withdrawal severity was associated with opioid physical dependence, linear regressions assessing associations between the number of somatic withdrawal signs expressed and total number of self-administered maintenance dose (12.5 μg/0.05 mL) heroin infusions were conducted. For data from Phase 2 not graphically depicted, means and standard deviations (SDs) for incidence of withdrawal signs are represented in Table 1. Note that Phase 2 data are comprised of a subset of Phase 1 animals and that data on the acute withdrawal syndrome from those animals are described under both phases. This was necessary to provide appropriate content for interpreting the protracted withdrawal symptoms. All analyses were conducted in GraphPad Prism 8.0.

**Table 1.** Mean Number Signs at Each Timepoint

|  | Baseline |  | 0 hr |  | 16 hrs |  | 48 hrs |  | 72 hrs |  | 168 hrs |  | 312 hrs |  |
| --- | --- | --- | --- | --- | --- | --- | --- | --- | --- | --- | --- | --- | --- | --- |
|  | Mean | SD | Mean | SD | Mean | SD | Mean | SD | Mean | SD | Mean | SD | Mean | SD |
| **Total Sample (N=16)** |  |  |  |  |  |  |  |  |  |  |  |  |  |  |
| Body Temperature (C°) | 34.6 | 1.3 | 34.6 | 1.2 | 34.7 | 1.3 | 35.1 | 1.6 | 35.2 | 1.3 | 35.3 | 1.5 | 36.4 | 1.0 |
| Body Shakes | 0.0 | 0.0 | 0.0 | 0.0 | 0.1 | 0.3 | 0.0 | 0.0 | 0.1 | 0.3 | 0.1 | 0.3 | 0.1 | 0.5 |
| Chews | 1.9 | 1.9 | 2.2 | 3.9 | 1.6 | 1.7 | 1.1 | 0.9 | 0.9 | 1.3 | 0.8 | 1.2 | 1.1 | 1.4 |
| Diarrhea/Defecation | 0.0 | 0.0 | 0.0 | 0.0 | 0.1 | 0.3 | 0.0 | 0.0 | 0.0 | 0.0 | 0.0 | 0.0 | 0.0 | 0.0 |
| Digs | 4.4 | 5.7 | 4.6 | 7.2 | 1.9 | 3.4 | 1.1 | 2.3 | 1.4 | 4.0 | 2.9 | 4.6 | 1.3 | 2.0 |
| Escape Attempts | 26.0 | 7.8 | 21.5 | 11.3 | 15.6 | 8.1 | 19.8 | 15.0 | 16.1 | 6.6 | 16.8 | 10.1 | 16.3 | 11.1 |
| Eye Blinks | 0.0 | 0.0 | 0.0 | 0.0 | 0.0 | 0.0 | 0.0 | 0.0 | 0.0 | 0.0 | 0.0 | 0.0 | 0.0 | 0.0 |
| Foot Licks | 4.0 | 2.9 | 4.4 | 4.3 | 19.1 | 16.1 | 28.2 | 32.7 | 17.3 | 16.9 | 16.3 | 20.0 | 25.5 | 35.5 |
| Genital Licks | 1.3 | 1.7 | 0.4 | 0.9 | 1.1 | 1.1 | 1.1 | 1.5 | 0.9 | 1.2 | 0.5 | 0.8 | 0.7 | 0.6 |
| Grooms | 5.8 | 2.7 | 7.5 | 6.5 | 18.7 | 7.8 | 9.3 | 7.0 | 7.1 | 4.8 | 9.1 | 5.3 | 7.2 | 5.2 |
| Head Shakes | 0.0 | 0.0 | 0.0 | 0.0 | 0.0 | 0.0 | 0.0 | 0.0 | 0.0 | 0.0 | 0.0 | 0.0 | 0.0 | 0.0 |
| Jumps | 1.6 | 3.1 | 1.2 | 2.5 | 0.4 | 1.0 | 0.4 | 0.6 | 0.6 | 1.6 | 1.1 | 2.2 | 0.8 | 2.5 |
| Piloerection | 0.0 | 0.0 | 0.0 | 0.0 | 0.0 | 0.0 | 0.0 | 0.0 | 0.0 | 0.0 | 0.0 | 0.0 | 0.0 | 0.0 |
| Ptois | 0.0 | 0.0 | 0.0 | 0.0 | 0.0 | 0.0 | 0.0 | 0.0 | 0.0 | 0.0 | 0.0 | 0.0 | 0.0 | 0.0 |
| Rears | 4.8 | 3.5 | 3.9 | 4.3 | 4.3 | 2.3 | 5.7 | 3.4 | 5.5 | 3.8 | 4.4 | 4.6 | 4.6 | 4.8 |
| Teeth Chatters | 0.0 | 0.0 | 0.0 | 0.0 | 0.1 | 0.3 | 0.1 | 0.3 | 0.1 | 0.3 | 0.0 | 0.0 | 0.0 | 0.0 |
| Writhes | 0.1 | 0.5 | 1.6 | 3.8 | 1.1 | 2.7 | 0.8 | 2.8 | 1.6 | 4.5 | 0.5 | 1.3 | 0.7 | 2.5 |
| **Total Somatic Signs** | **49.9** | **29.8** | **47.4** | **44.6** | **64.1** | **45.2** | **67.4** | **66.5** | **51.8** | **45.1** | **52.4** | **50.4** | **58.1** | **66.1** |
| **Male (N=8)** |  |  |  |  |  |  |  |  |  |  |  |  |  |  |
| Body Temperature (C°) | 34.8 | 1.4 | 34.1 | 1.0 | 34.0 | 0.7 | 34.7 | 1.0 | 34.3 | 0.8 | 34.8 | 0.8 | 36.0 | 1.1 |
| Body Shakes | 0.0 | 0.0 | 0.0 | 0.0 | 0.0 | 0.0 | 0.0 | 0.0 | 0.1 | 0.4 | 0.0 | 0.0 | 0.3 | 0.7 |
| Chews | 2.1 | 1.5 | 3.6 | 5.2 | 2.5 | 1.9 | 1.0 | 0.9 | 1.5 | 1.6 | 1.4 | 1.3 | 1.3 | 1.2 |
| Diarrhea/Defecation | 0.0 | 0.0 | 0.0 | 0.0 | 0.0 | 0.0 | 0.0 | 0.0 | 0.0 | 0.0 | 0.0 | 0.0 | 0.0 | 0.0 |
| Digs | 2.3 | 2.6 | 2.3 | 3.5 | 0.9 | 1.5 | 0.0 | 0.0 | 0.1 | 0.4 | 1.1 | 2.1 | 0.6 | 0.7 |
| Escape Attempts | 26.6 | 6.9 | 24.8 | 12.7 | 11.8 | 5.9 | 14.1 | 17.0 | 14.8 | 6.5 | 16.4 | 11.6 | 13.6 | 10.6 |
| Eye Blinks | 0.0 | 0.0 | 0.0 | 0.0 | 0.0 | 0.0 | 0.0 | 0.0 | 0.0 | 0.0 | 0.0 | 0.0 | 0.0 | 0.0 |
| Foot Licks | 5.4 | 3.3 | 5.5 | 3.0 | 14.1 | 5.0 | 18.1 | 18.0 | 12.5 | 9.7 | 9.5 | 6.4 | 18.8 | 31.6 |
| Genital Licks | 1.5 | 2.0 | 0.3 | 0.5 | 1.1 | 1.1 | 1.3 | 2.1 | 1.0 | 1.4 | 0.6 | 1.1 | 0.9 | 0.6 |
| Grooms | 6.3 | 3.0 | 5.8 | 3.4 | 20.8 | 4.6 | 7.0 | 4.8 | 7.5 | 5.9 | 9.6 | 5.3 | 4.9 | 3.0 |
| Head Shakes | 0.0 | 0.0 | 0.0 | 0.0 | 0.0 | 0.0 | 0.0 | 0.0 | 0.0 | 0.0 | 0.0 | 0.0 | 0.0 | 0.0 |
| Jumps | 0.4 | 0.7 | 1.6 | 3.1 | 0.0 | 0.0 | 0.3 | 0.7 | 1.1 | 2.2 | 1.5 | 2.8 | 1.5 | 3.5 |
| Piloerection | 0.0 | 0.0 | 0.0 | 0.0 | 0.0 | 0.0 | 0.0 | 0.0 | 0.0 | 0.0 | 0.0 | 0.0 | 0.0 | 0.0 |
| Ptois | 0.0 | 0.0 | 0.0 | 0.0 | 0.0 | 0.0 | 0.0 | 0.0 | 0.0 | 0.0 | 0.0 | 0.0 | 0.0 | 0.0 |
| Rears | 5.1 | 3.8 | 4.5 | 5.6 | 4.4 | 2.3 | 6.1 | 3.8 | 5.3 | 4.3 | 5.9 | 5.9 | 6.1 | 6.1 |
| Teeth Chatters | 0.0 | 0.0 | 0.0 | 0.0 | 0.3 | 0.5 | 0.1 | 0.4 | 0.1 | 0.4 | 0.0 | 0.0 | 0.0 | 0.0 |
| Writhes | 0.0 | 0.0 | 0.4 | 1.1 | 0.5 | 1.4 | 0.0 | 0.0 | 0.0 | 0.0 | 0.1 | 0.4 | 0.1 | 0.4 |
| **Total Somatic Signs** | **49.6** | **23.8** | **48.6** | **38.0** | **56.3** | **24.1** | **48.0** | **47.7** | **44.0** | **32.7** | **46.1** | **36.7** | **48.0** | **58.4** |
| **Female (N=8)** |  |  |  |  |  |  |  |  |  |  |  |  |  |  |
| Body Temperature (C°) | 34.3 | 1.3 | 35.1 | 1.2 | 35.4 | 1.3 | 35.5 | 2.0 | 36.1 | 1.0 | 35.9 | 1.8 | 36.9 | 0.6 |
| Body Shakes | 0.0 | 0.0 | 0.0 | 0.0 | 0.1 | 0.4 | 0.0 | 0.0 | 0.0 | 0.0 | 0.1 | 0.4 | 0.0 | 0.0 |
| Chews | 1.6 | 2.4 | 0.8 | 1.2 | 0.8 | 1.0 | 1.1 | 1.0 | 0.4 | 0.5 | 0.3 | 0.7 | 1.0 | 1.7 |
| Diarrhea/Defecation | 0.0 | 0.0 | 0.0 | 0.0 | 0.1 | 0.4 | 0.0 | 0.0 | 0.0 | 0.0 | 0.0 | 0.0 | 0.0 | 0.0 |
| Digs | 6.5 | 7.2 | 7.0 | 9.2 | 3.0 | 4.4 | 2.1 | 2.9 | 2.8 | 5.5 | 4.6 | 5.9 | 1.9 | 2.7 |
| Escape Attempts | 25.4 | 9.1 | 18.3 | 9.3 | 19.4 | 8.5 | 25.5 | 10.9 | 17.5 | 6.9 | 17.1 | 9.3 | 18.9 | 11.7 |
| Eye Blinks | 0.0 | 0.0 | 0.0 | 0.0 | 0.0 | 0.0 | 0.0 | 0.0 | 0.0 | 0.0 | 0.0 | 0.0 | 0.0 | 0.0 |
| Foot Licks | 2.6 | 1.8 | 3.4 | 5.2 | 24.1 | 21.8 | 38.3 | 41.6 | 22.1 | 21.5 | 23.1 | 26.7 | 32.3 | 39.9 |
| Genital Licks | 1.1 | 1.5 | 0.6 | 1.2 | 1.0 | 1.2 | 1.0 | 0.8 | 0.9 | 1.0 | 0.4 | 0.5 | 0.5 | 0.5 |
| Grooms | 5.3 | 2.4 | 9.3 | 8.5 | 16.6 | 10.1 | 11.5 | 8.4 | 6.8 | 3.8 | 8.5 | 5.6 | 9.5 | 6.1 |
| Head Shakes | 0.0 | 0.0 | 0.0 | 0.0 | 0.0 | 0.0 | 0.0 | 0.0 | 0.0 | 0.0 | 0.0 | 0.0 | 0.0 | 0.0 |
| Jumps | 2.9 | 4.1 | 0.8 | 1.8 | 0.8 | 1.4 | 0.5 | 0.5 | 0.1 | 0.4 | 0.8 | 1.5 | 0.0 | 0.0 |
| Piloerection | 0.0 | 0.0 | 0.0 | 0.0 | 0.0 | 0.0 | 0.0 | 0.0 | 0.0 | 0.0 | 0.0 | 0.0 | 0.0 | 0.0 |
| Ptois | 0.0 | 0.0 | 0.0 | 0.0 | 0.0 | 0.0 | 0.0 | 0.0 | 0.0 | 0.0 | 0.0 | 0.0 | 0.0 | 0.0 |
| Rears | 4.5 | 3.4 | 3.4 | 2.8 | 4.3 | 2.5 | 5.3 | 3.2 | 5.8 | 3.5 | 2.9 | 2.3 | 3.0 | 2.6 |
| Teeth Chatters | 0.0 | 0.0 | 0.0 | 0.0 | 0.0 | 0.0 | 0.0 | 0.0 | 0.0 | 0.0 | 0.0 | 0.0 | 0.0 | 0.0 |
| Writhes | 0.3 | 0.7 | 2.8 | 5.1 | 1.8 | 3.6 | 1.6 | 3.9 | 3.3 | 6.1 | 0.9 | 1.8 | 1.3 | 3.5 |
| **Total Somatic Signs** | **50.1** | **32.5** | **46.1** | **44.3** | **71.9** | **55.2** | **86.9** | **73.2** | **59.5** | **49.1** | **58.6** | **54.6** | **68.3** | **68.8** |

Data represent absolute count of withdrawal symptoms following abrupt opioid discontinuation. Values are the number of occurrences observed by raters blinded to the timepoint during a 10-minute observation period (following a 5-minute habituation) for each symptom. Timepoint represents hours since last heroin self-administration. Data represent values from Experiment 2 only.

## Results

### Heroin Self-Administration

#### Overall

Results from a three-way ANOVA revealed a significant main effect of sex (F_1,392_ = 10.54, *p* < 0.01), and lever (F_1,392_ = 87.79, *p* < 0.0001), and a sex x lever interaction (F_1,392_ = 3.86, *p* = 0.05). No main effects of session, session x lever, or session x sex x lever interaction were found (*p*’s>0.05). A one-way repeated measures ANOVA revealed that animals earned significantly more infusions over sessions (F_13,370_ = 7.12, *p* < 0.0001; Figure 1A).

**Figure 1.**
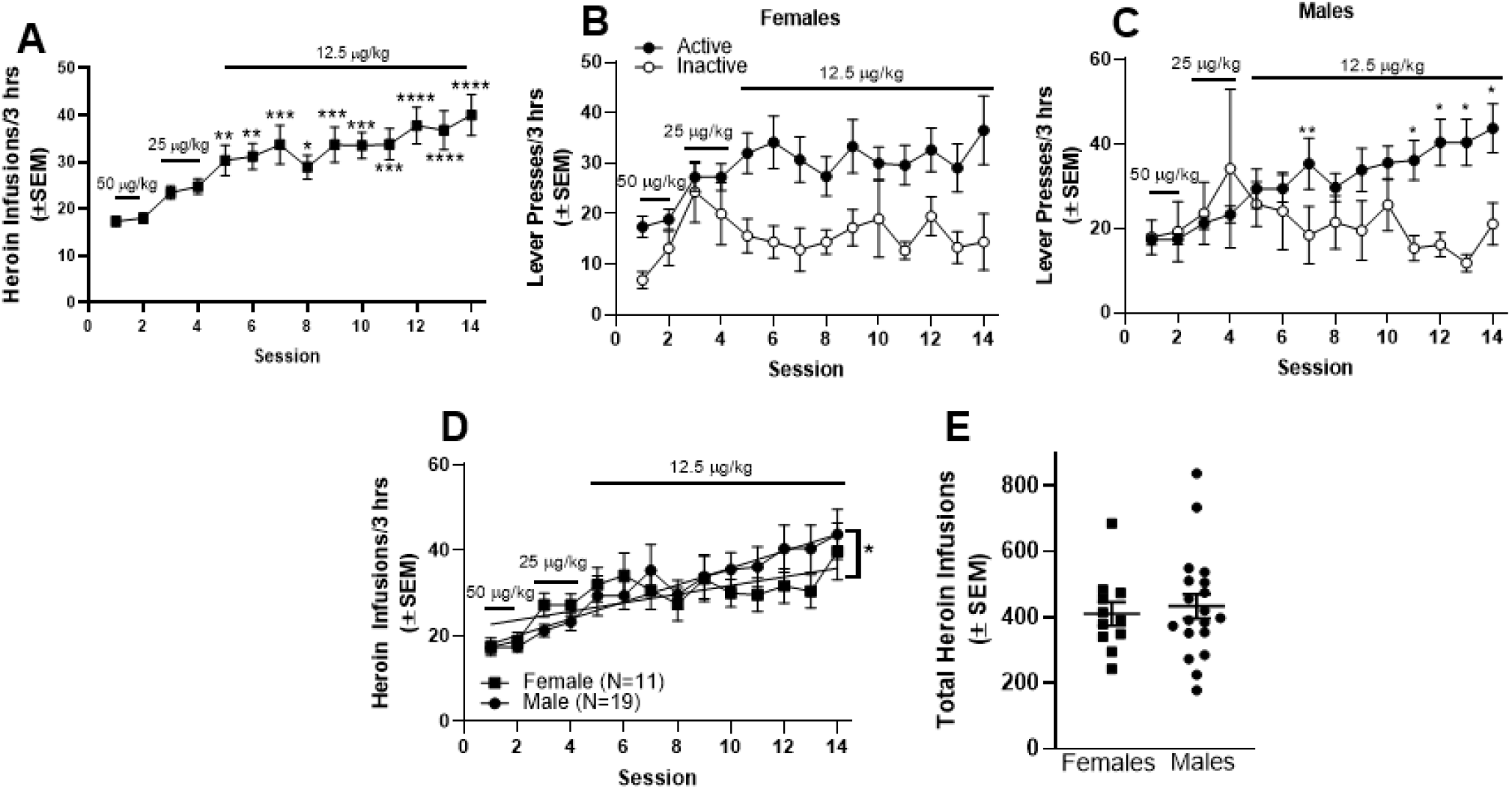
Heroin Self-Administration. **(A)** Male and female rats showed increased heroin intake across sessions, comparing infusions earned in each session versus session 1. When broken down by sex, both females **(B)** and males **(C)** showed discrimination between the active and inactive levers across sessions. **(D)** The rate of heroin intake increased significantly faster in males than in females. **(E)** Total number of heroin infusions across all 14 self-administration sessions did not differ as a function of sex. In (A): \**p*<0.05, \*\**p*<0.01, \*\*\**p*<0.001, \*\*\*\**p*<0.0001 compared to session 1. In (C): \**p*<0.05, \*\**p*<0.01 comparing active to inactive lever pressing. In (D): \**p*<0.05 comparing the slopes of male and female heroin intake fitted curves.

#### Sex-Specific Effects

Data suggest that males and females discriminated between the active and inactive levers. Female rats demonstrated significant main effects of session (F_13,280_ = 2.49, *p* < 0.01) and lever (F_1,20_ = 16.65, *p* < 0.001) on self-administration though no significant lever x session interaction was observed (*p* > 0.05; Figure 1B). An analysis of active and inactive lever pressing in male rats demonstrated significant main effects of session (F_13,468_ = 1.71, *p* < 0.05), lever (F_1,36_ = 9.35, *p* < 0.01), and a significant session x lever interaction (F_13,468_ = 2.40, *p* < 0.01; Figure 1C) that was driven by differences in active and inactive lever presses on sessions 7 and 11-14.

In addition, although both males and females demonstrated a significant effect of sessions on number of infusions over time, the rate of intake was faster in males versus females. A two-way repeated measures ANOVA with sex as between and session as within-subjects factors revealed a significant main effect of session (F_13,358_ = 6.66, *p* < 0.0001), but no main effect of sex or sex x session interaction. Linear regressions conducted to further determine if sex differences in intake occurred across sessions revealed the slope of the acquisition curve was significantly different from zero in both males (F_1,264_ = 51.41, *p* < 0.0001) and females (F_1,152_ = 13.98, *p* < 0.001). The slopes of the male and female infusion curves were also significantly different from each other (F_1,416_ = 5.38, *p* < 0.05) in a manner that suggested the rate of heroin intake increased faster in males compared to females (see Figure 1D). Total number of heroin infusions revealed no significant difference in total heroin intake as a function of sex (*t*(28) = 0.42, *p* > 0.05; Figure 1E).

### Phase 1: Defining Acute Heroin Withdrawal following Heroin Self-Administration

#### Total Somatic Signs of Acute Heroin Withdrawal

##### Overall

A timeline of experimental procedures is shown in Figure 2A and the results showed that animals experienced expected increases in some symptoms following discontinuation of heroin self-administration. Overall, the total number of spontaneous somatic signs significantly differed across the 4 evaluated timepoints (0, 16, 48, 72 hrs; F_(3, 87)_=5.44, *p*<0.01) (Figure 2B). Post-hoc testing revealed that total number of symptoms increased significantly between the 0 hr timepoint and the 16 and 48 hr timepoints.

**Figure 2.**
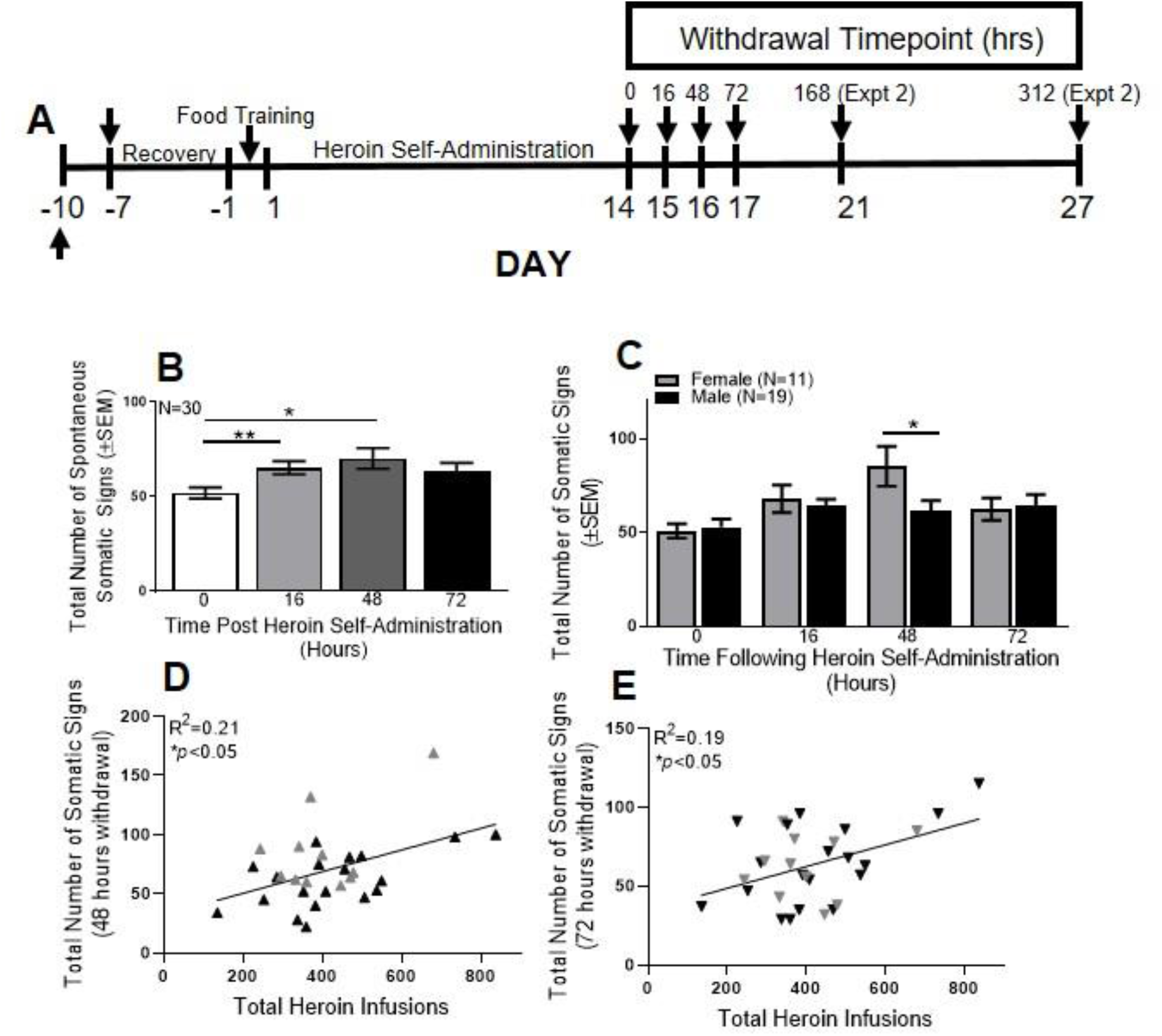
Total Somatic Signs during the Acute Withdrawal Period. **(A)** Experimental timeline for Phases 1 and 2. Withdrawal timepoints were taken for 0, 16, 48, and 72 hrs (Phase 1) following heroin self-administration, and at baseline, 0, 16, 48, 72, 168, and 312 hrs (Phase 2). **(B)** Total number of somatic signs (excluding body temperature) increased from 0 to 48 hrs following cessation of heroin self-administration. Total number of somatic signs positively correlated with heroin intake at the **(C)** 48 and **(D)** 72 hr timepoints. Gray triangles indicate females; black triangles indicate males. **(E)** Females showed significantly elevated total somatic signs of withdrawal compared to males at the 48 hr timepoint. In B: \*\**p*<0.01 compared to 0 hrs; \**p*<0.05 compared to 0 hrs. In E: \**p*<0.05 comparing males vs. females. Data are represented as mean ± standard error of the mean (SEM).

##### Sex-Specific Effects

A significant main effect of timepoint (F_(3,84)_ = 7.37, *p*<0.001) and a significant sex x timepoint interaction (F_(3,84)_ = 3.33, *p*<0.05) on total withdrawal were also observed. Post-hoc testing indicated this was driven by females expressing significantly more somatic signs of withdrawal at the 48 hr timepoint than males (Figure 2C).

##### Relationship to Heroin Self-administration

Linear regressions found no significant association between somatic withdrawal symptoms and the total number of heroin infusions during the self-administration period between the 0 and 16 hrs timepoints (data not shown; *p*’s>0.05), but significant positive correlations were observed for the 48 hr (F_(1,28)_=7.38, *p*<0.05, R^2^=0.21; Figure 2D) and 72 hr (F_(1,28)_=7.38, *p*<0.05, R^2^=0.21; Figure 2E) timepoints.

#### Anxiety/Escape Behaviors

##### Overall

Animals showed minimal evidence of symptoms classified as representing anxiety. Specifically, there was no significant effect of digging (*p*>0.05; Figure 3A), jumping (*p*>0.05; Figure 3B), or rearing (*p*>0.05; Figure 3C) across the 4 timepoints. A significant effect of time was observed for escape attempts (F_(3, 87)_=3.74, *p*<0.05; Figure 3D), which post-hoc comparisons revealed was driven by a significant decrease in escape attempts between the 16 and 48 hr timepoints.

**Figure 3.**
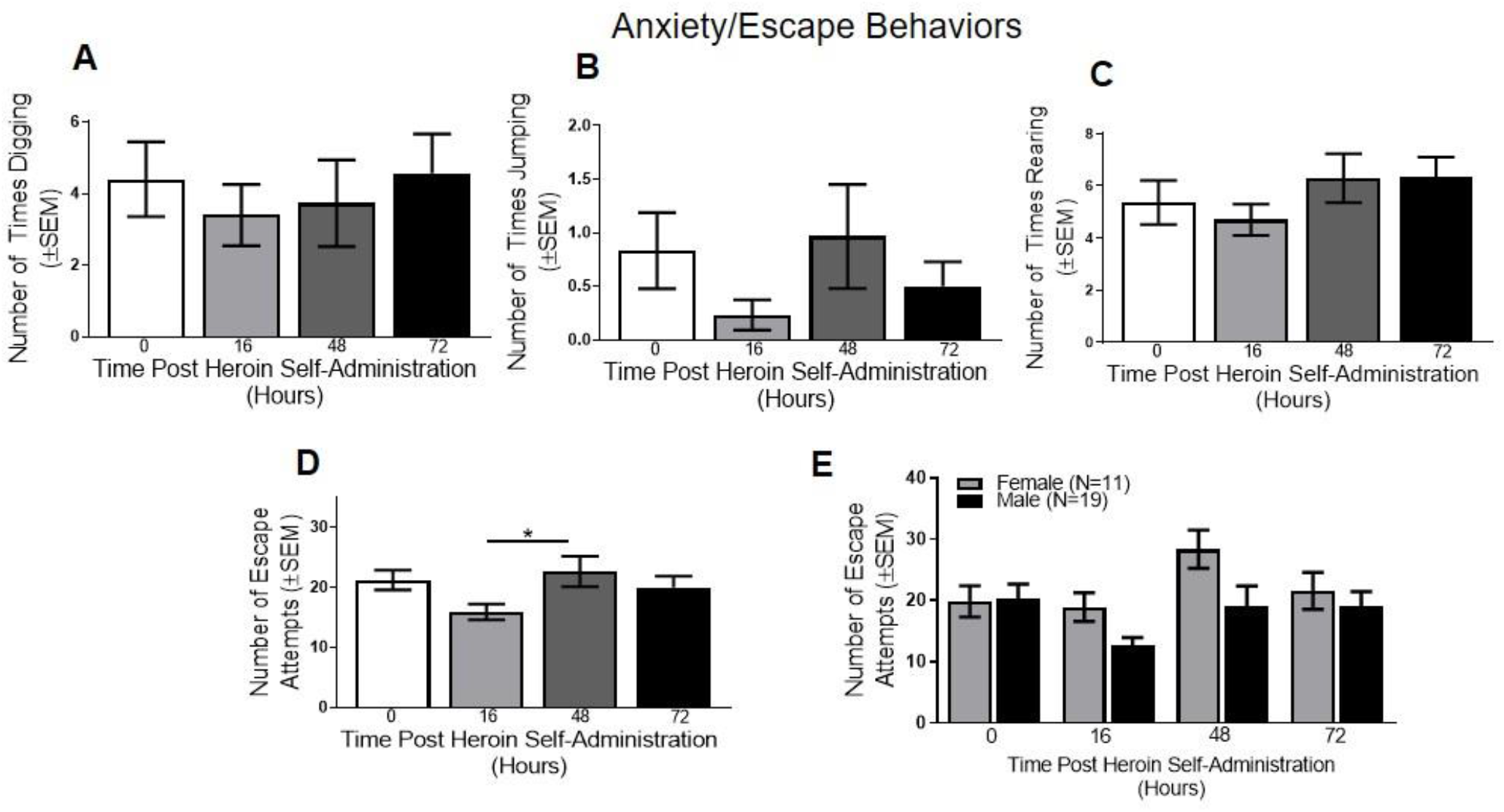
Anxiety/Escape Behaviors in the Acute Withdrawal Period. **(A)** Digging, **(B)** jumping, **(C)** and rearing did not differ as a function of time following heroin self-administration. **(D)** Escape behaviors varied as a function of time following heroin self-administration, with escape attempts increasing between 16 and 48 hrs. **(E)** When broken down by sex, a significant main effect of timepoint was found. In D: \**p*<0.05 comparing 16 to 48 hr withdrawal timepoints. Data are represented as mean ± SEM.

##### Sex-Specific Effects

No significant main effects of sex or timepoint or sex x timepoint interactions were observed for digging, jumping, or rearing (*p*’s>0.05; data not shown). A significant main effect of timepoint was found in escape attempts (F_(3, 84)_=4.61, *p*<0.01; 3E), though the main effect of sex or sex x timepoint interaction were not significant (*p*’s>0.05).

#### Body Temperature

##### Overall

No significant effect of body temperature (*p*>0.05; Figure 4A) was observed across the 4 timepoints when sex was collapsed. Although the main effect of time not reach significance (*p*>0.05), there were notable variations in teeth chattering across the timepoints (see Figure 4B).

**Figure 4.**
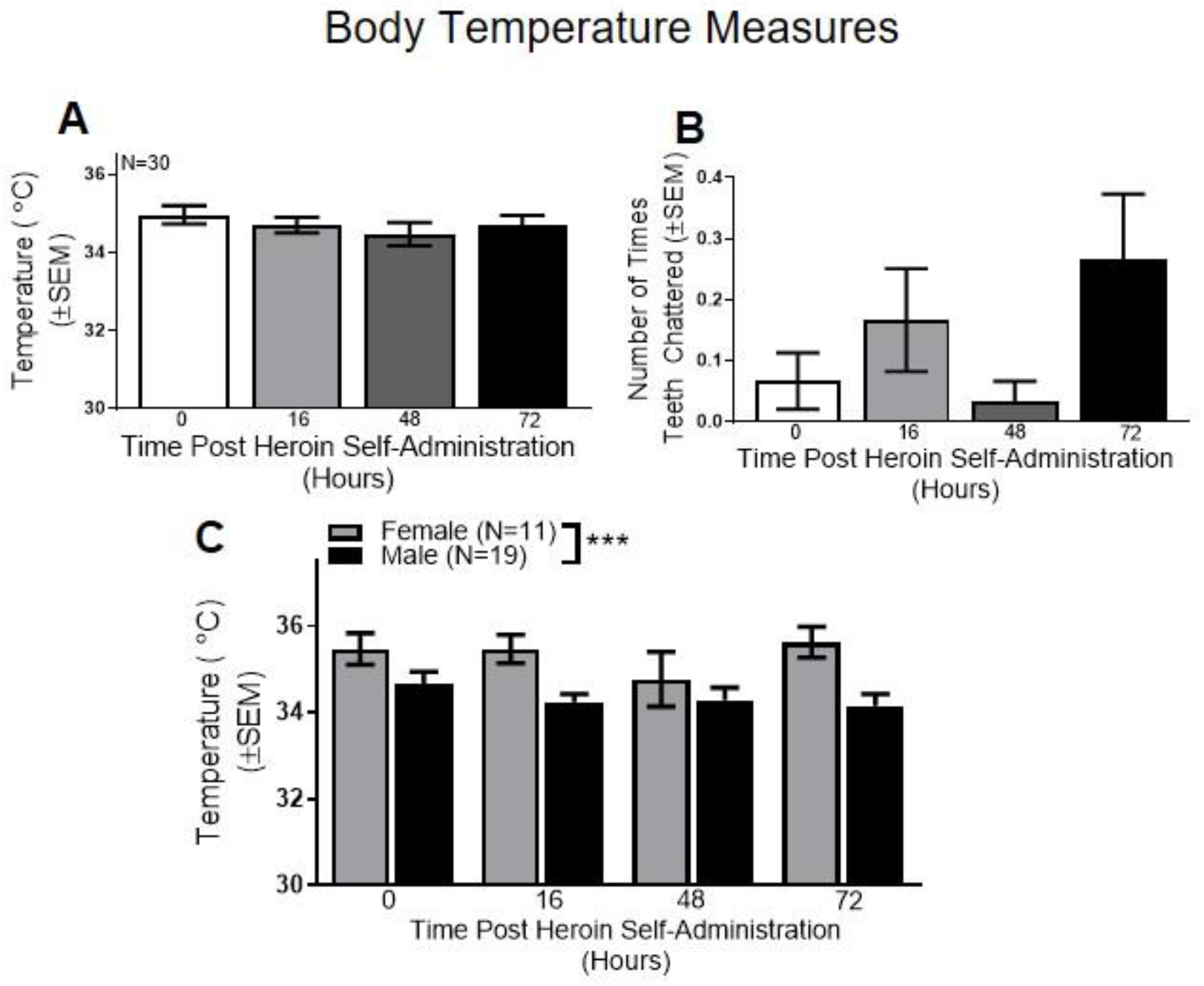
Body Temperature Measures in the Acute Withdrawal Period. Overall, no differences were found across the 4 acute timepoints with **(A)** body temperature or **(B)** teeth chattering. **(C)** When broken down by sex, a significant main effect of sex was found in body temperature. \*\*\**p*<0.001 comparing males vs. females. Data are represented as mean ± SEM.

##### Sex-Specific Effects

Sex-specific analysis of body temperature revealed a significant main effect of sex (F_(1,28)_=21.32, *p*>0.0001; Figure 4C), but no main effect of time or sex x time interaction (*p*’s>0.05). These results indicate that females had significantly higher body temperature compared to males. No significant main effects of sex or time or sex x time interaction was found for teeth chattering (*p*’s>0.05; data not shown).

#### Gastrointestinal Symptoms

##### Overall

No significant change in diarrhea (*p*>0.05) or defecation (*p*>0.05) were observed at any timepoint, with only one animal showing signs of diarrhea at one timepoint (data not shown).

##### Sex-Specific Effects

No sex-related significant effects were observed for diarrhea or defecation (*p*’s>0.05).

#### Hyperactivity

##### Overall

Several symptoms within the category of hyperactivity had significant effects and/or displayed meaningful trends. For instance, a significant effect of grooming was observed (F_(3, 87)_=18.00, *p*<0.0001; Figure 5A) that was driven by a significant difference between the 0 and 16 hr, 16 and 48 hr, and 16 and 72 hr timepoints, indicating that grooming peaks within one day of heroin withdrawal and declines thereafter. A significant effect of foot licking was also found (F_(3, 55)_=3.13, *p*<0.05; Figure 5B), which was driven by a significant increase in foot licking at 16 hrs relative to all other timepoints, concurrent with the increase in grooming. No significant effects of chewing (*p*>0.05, Figure 5C) or genital licking (*p*>0.05, Figure 5D) were observed.

**Figure 5.**
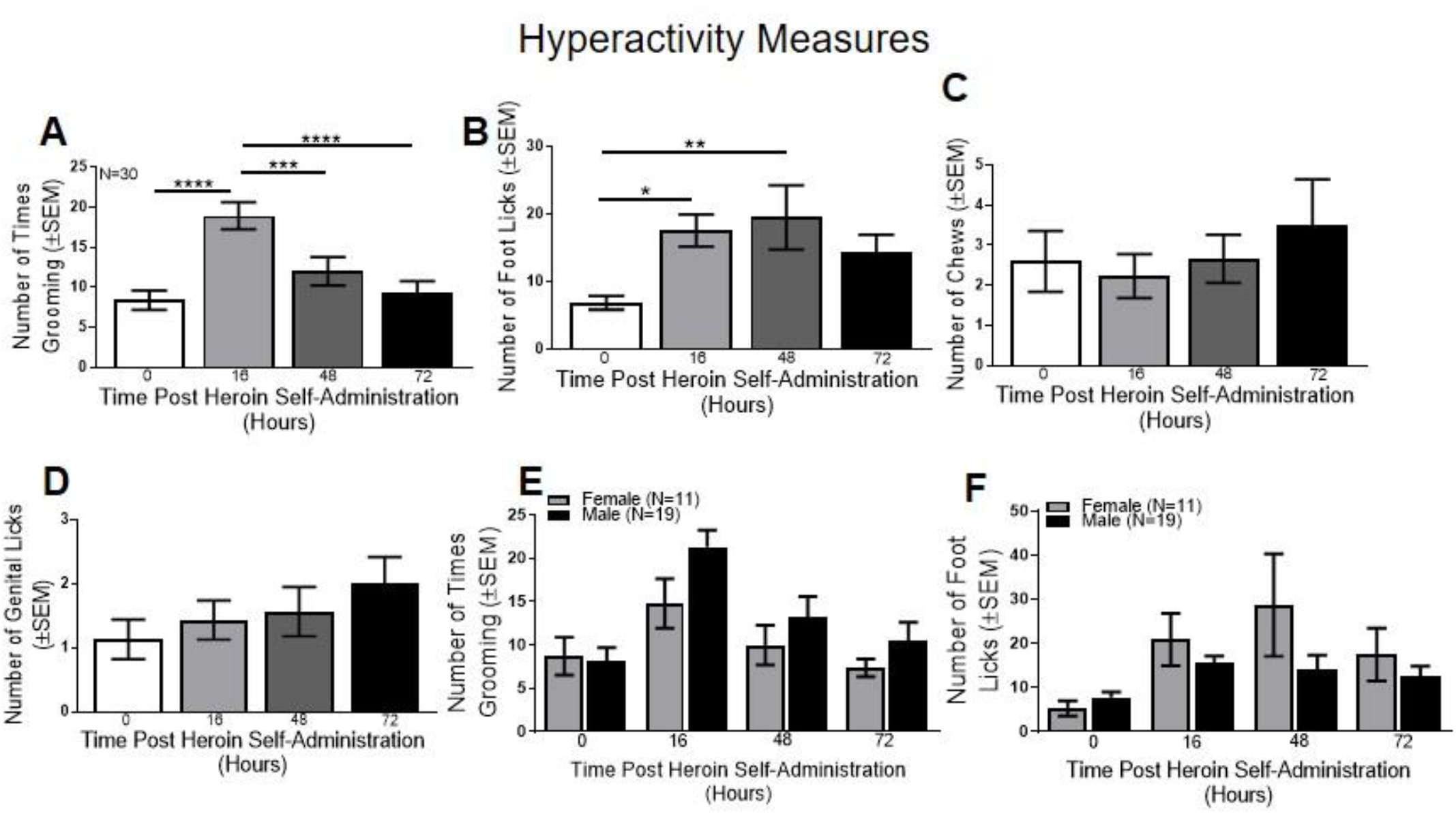
Hyperactivity Measures in the Acute Withdrawal Period. **(A)** Overall, number of times grooming increased significantly between the 0 and 16 hr timepoints, and then began to decrease by 48 hrs. **(B)** Number of footlicks also increased between the 0 and 16 hr timepoints, but remained elevated through the 48 hr timepoint before decreasing at 72 hrs. Number of **(C)** chews and **(D)** genital licks did not differ as a function of withdrawal timepoint. **(E)** When broken down by sex, there was a significant main effect of timepoint for grooming, indicating that grooming increased by the 16 hr timepoint in an non-sex-dependent manner. **(F)** Number of footlicks also differed as a function of time, but not as function of sex. In A: \*\*\*\**p*<0.0001 comparing 0 to 16 hrs or 16 to 72 hrs; \*\*\**p*<0.001 comparing 16 to 48 hrs. In B: \**p*<0.05 comparing 0 to 16 hrs; \*\**p*<0.01 comparing 0 to 48 hrs. Data are represented as mean ± SEM.

##### Sex-Specific Effects

A significant main effect of timepoint was found for grooming (F_(3, 84)_=14.70, *p*<0.0001; Figure 5E), but the main effect of sex and sex x timepoint interactions were not significant (*p*’s>0.05). A significant main effect of timepoint on footlicks was also identified (F_(3, 84)_=6.42, *p*<0.001; Figure 5F), with no main effect of sex or sex x timepoint interactions. Finally, genital licking revealed no main effects of sex or timepoint or sex x timepoint interactions (*p*’s>0.05, data not shown).

#### Pain, Hyperalgesia and Ptosis

##### Overall

No significant change in writhing or ptosis (*p*’s>0.05; data not shown) were observed.

##### Sex-Specific Effects

A significant main effect of timepoint was observed for writhing (F_(3, 75)_=4.81, *p*<0.01) and the main effect of sex trended but did not reach significance (F_(1,25)_=3.61, *p*=0.06). The sex x timepoint interaction was not significant (*p*>0.05). Inspection of the data suggested that both sexes evidenced a decline in writhing over days and that females demonstrated a higher number of writhes than males in each timepoint (data not shown). Ptosis revealed no main effects or interactions (*p*’*s*>0.05, data not shown).

#### Tremors/Shaking

##### Overall

No main effects or interactions were observed for head shakes (*p*>0.05; data not shown) or body shakes (*p*>0.05; data not shown) across the four timepoints.

##### Sex-Specific Effects

No main effects of sex or interactions were observed for head shakes or body shakes (*p*’*s*>0.05, data not shown).

### Phase 2: Defining a Preclinical Protracted Heroin Withdrawal Syndrome Following Self-Administration

#### Total Somatic Signs of Acute and Protracted Withdrawal

##### Overall

In contrast to Phase 1, once the baseline and 1- and 2-week timepoints were included in the model, the significant effect of timepoint (0, 16, 48, 72, 168, and 312 hrs) on somatic signs of withdrawal trended but no longer met statistical significance (F_(6,90)_=2.1, *p*=0.06; Figure 6A). However, a notable increase in symptom expression was observed at the 16 and 48 hr timepoints. Symptom counts for each time point are presented in Table 1 as a for the overall group as well as male and female subsamples.

**Figure 6.**
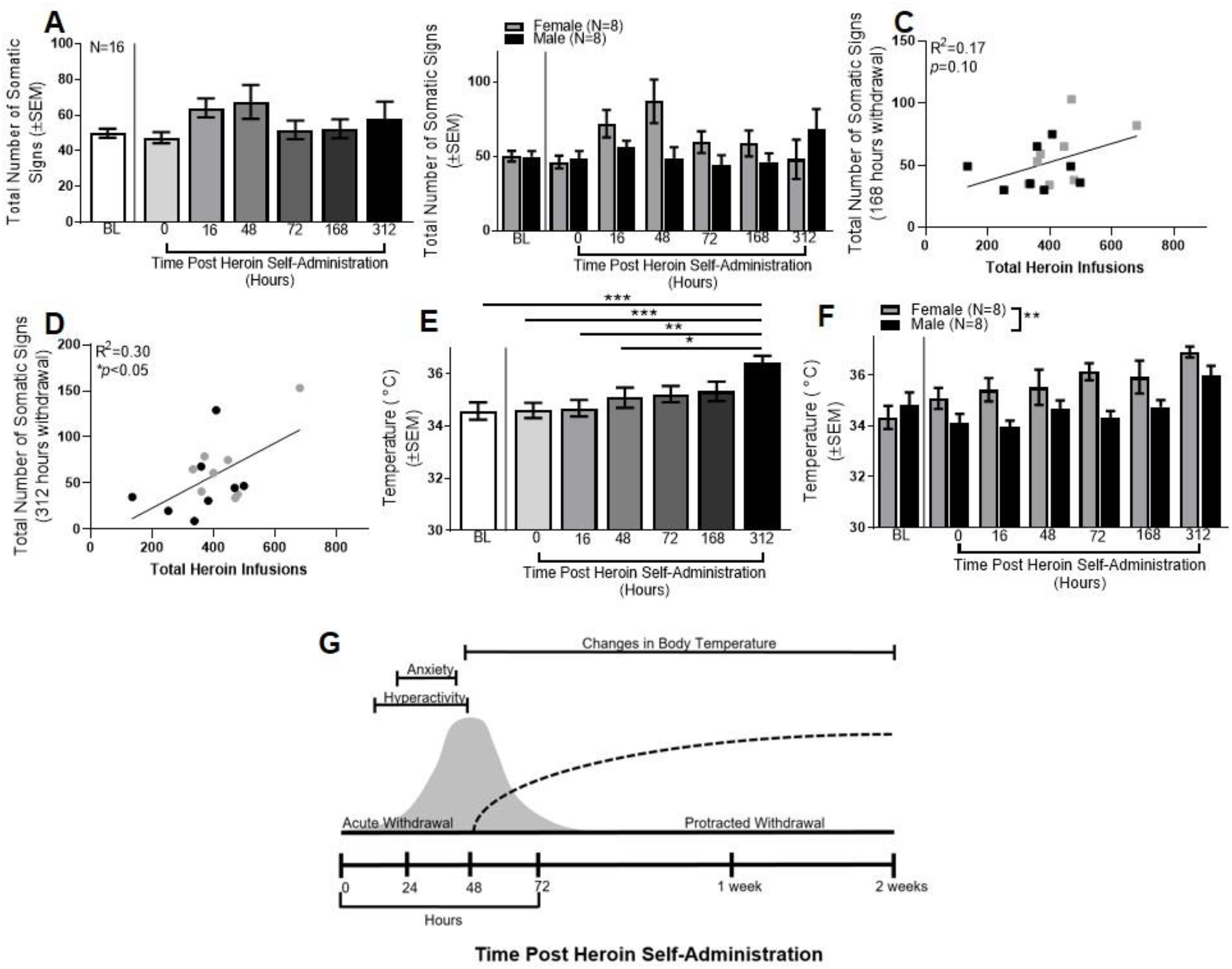
Defining the Protracted Heroin Withdrawal Syndrome Following Self-Administration. **(A)** Overall, total number of somatic signs did not differ as a function of time when baseline and 6 withdrawal timepoints were assessed. **(B)** When total number of somatic signs was broken down by sex, a significant effect of timepoint emerged. **(C)** Total number of somatic signs did not correlate with total number of heroin infusions at the 168 hr timepoint. Gray squares indicate females, black squares indicate males. **(D)** At 312 hrs, a significant positive correlation was found between total number of somatic signs and total heroin infusions. Gray circles indicate females, black circles indicate males. **(E)**Overall, a significant effect of timepoint was observed for body temperature, which was driven by significant differences between baseline and 312 hrs, as well as 0 and 312, 16 and 312, and 48 versus 312 hrs. **(F)** When broken down by sex, significant main effects of sex and timepoint were found. **(G)** Schematic representation of the acute and protracted withdrawal periods following heroin self-administration in rats. In E: \*\*\**p*<0.001 comparing baseline or 0 to 312 hrs; \*\**p*<0.01 comparing 16 to 312 hrs. \**p*<0.05 comparing 48 to 312 hrs. Data are represented as mean ± SEM.

##### Sex-Specific Effects

Sex-specific analyses revealed a significant main effect of timepoint (F_(6,84)_=2.21, *p*=0.05; Figure 6B), although no main effect of sex or sex x timepoint interaction was found (*p*’s>0.05). Though it did not reach the level of significance, inspection of data revealed that withdrawal expression was higher among females relative to males at the 16 and 48 hr timepoints.

##### Relationship to Heroin Self-Administration

The correlation between heroin infusions and total somatic signs did not reach significance at the 168 hr timepoint (F_(1,14)_=2.98, *p*>0.05, R^2^=0.17; Figure 6C), but did reach significance at the 312 hr timepoint (F_(1,14)_=6.10, *p*<0.05, R^2^=0.30; Figure 6D). This latter finding was surprising since the total number of signs at the protracted timepoints did not vary significantly from baseline (see Figure 6A). The fact that withdrawal at this timepoint was related to the number of heroin infusions at baseline suggests that self-administration was sufficient to produce a protracted opioid withdrawal syndrome.

#### Anxiety/Escape Behaviors

##### Overall

A significant effect was observed for digging (F_(6,90)_=3.19, *p*<0.05), with digging instances decreasing across time. As well, a significant main effect of timepoint for escape attempts was found (F_(6,90)_=3.41, *p*<0.01), with attempts decreasing across time. There was no significant effect of rearing or jumping (*p*’s>0.05).

##### Sex-Specific Effects

There was a significant sex x timepoint interaction on jumping (F_(6,84)_=2.29, *p*<0.05), which post-hoc testing revealed was driven by differences between the baseline and 312 hour timepoints in females (data not shown). There was no main effect of sex or sex x timepoint interactions observed for digging, escape attempts, or rearing (*p*’s>0.05).

#### Body Temperature

##### Overall

A significant effect of timepoint (F_(6,90)_=4.89, *p*<0.001; Figure 6E) was observed for body temperature, which was driven by significant differences between baseline and 312 hours, as well as 0 and 312, 16 and 312, and 48 versus 312 hours. These results indicate that body temperature increased after extended withdrawal (almost 2 weeks following heroin self-administration), further supporting the notion that a 312 hr rating may detect a protracted withdrawal syndrome. There were no main effects or interactions in the teeth chattering category (*p*’s>0.05).

##### Sex-specific Effects

Analyses revealed a significant main effect of timepoint (F_(6,84)_=5.07, *p*<0.001; Figure 6F) and sex (F_(1,14)_=13.28, *p*<0.01), but no sex x timepoint interaction on body temperature. When broken down by sex, there was no main effect of sex, timepoint or sex x timepoint interaction in the teeth chattering category (*p*’s>0.05).

#### Gastrointestinal Symptoms

##### Overall

No instances of diarrhea/defecation were observed for any of the additional timepoints (baseline, and 168 and 312 hours post heroin self-administration), so the main effects and interactions for diarrhea (*p*>0.05) or defecation (*p*>0.05) were not significant.

##### Sex-Specific Effects

No sex-related significant effects were observed for diarrhea or defecation (*p*’s>0.05).

#### Hyperactivity

##### Overall

A significant effect of timepoint on grooming was evident (F_(6,90)_=13.45, *p*<0.0001), with an increase by 16 hrs followed by a decrease. A significant effect was also observed for foot-licking (F_(6,90)_=4.19, *p*=0.001), with an increase by 16 hrs that remained elevated throughout the remaining timepoints. No effects were observed for chewing or genital licking (*p*’s>0.05).

##### Sex-specific Effects

A significant main effect of sex was observed for chewing (F_(1,14)_=5.91, *p*<0.05), but no significant main effect of timepoint or sex x timepoint interaction (*p*’s>0.05), suggesting that males had a significantly higher number of chews than females independent of timepoint. Chewing did not reveal a significant main effect of sex (*p*>0.05), suggesting that grooming behavior changed over time in a non-sex dependent way. No main effect of sex or sex x timepoint interaction was found for foot licking (*p*’s>0.05), indicating that foot licking differed across time in a non-sex-dependent manner. No sex-specific effects were found for genital licking (*p*’s>0.05).

#### Pain, Hyperalgesia and Ptosis

##### Overall

No significant change in writhing or ptosis (*p*’*s*>0.05) were observed.

##### Sex-Specific Effects

No significant main effects of sex or timepoint, or sex x timepoint interaction for writhing or ptosis were observed (*p*’*s*>0.05).

#### Tremors/Shaking

##### Overall

There were no instances of head shakes at any timepoint and no main effects or interactions for body shakes (*p*>0.05) across the 6 timepoints.

##### Sex-Specific Effects

No main effects of sex or interactions were observed for head shakes or body shakes (*p*’*s*>0.05).

## Discussion

The present study evaluated opioid withdrawal symptomatology over a 72 hr (Phase 1) and a 312 hr (Phase 2) period following a 14-day heroin self-administration and abrupt omission procedure in rats. This approach represents a novel investigation of the spontaneous heroin withdrawal syndrome following volitional heroin self-administration in male and female rats that, to our knowledge, is the first characterization of both the acute and protracted withdrawal syndrome in a preclinical rodent model of spontaneous opioid withdrawal. The signs selected for evaluation were those commonly reported in studies that administered opioids to animals and examined opioid withdrawal following naloxone administration (Dunn et al., 2019). In Phase 1, some (but not all) symptoms were evident within the 72 hr period. In Phase 2, several symptoms were still evident and even increasing in severity during the protracted monitoring period (see Table 1). In both cases, the rate of heroin self-administration was positively correlated with total overall symptom expression for many of the later timepoints, supporting validity of this model. As well, contrary to our hypothesis, heroin infusions increased significantly faster in male than female rats. However, both sexes acquired heroin self-administration.

To date, only one study has directly compared naloxone-precipitated and spontaneous opioid withdrawal in rats. That study found that precipitated withdrawal resulted in a more severe withdrawal syndrome than spontaneous withdrawal (Schulteis et al., 1998), and those results are consistent with the outcomes observed in this examination. Although some of the signs of spontaneous withdrawal evaluated in this study (e.g., escape attempts, body temperature, grooming, and foot licking) were evident over the initial 72-hr period, many others (e.g., diarrhea, defecation, writhing, ptosis, and head and body shakes) did not vary as a function of time and/or were never observed. Consistent with withdrawal in humans, symptoms also emerged at and were observable for different lengths of time. For instance, grooming and foot licking were the first symptoms to emerge (at the 16 hr timepoint) yet they lasted for different periods of time (grooming declined by 48 hours, whereas foot licking started declining at 72 hours). Several other symptoms emerged at the 16 hr timepoint and increased over time, including escape attempts and body temperature elevations.

This study also examined symptom expression during a protracted period (up to 312 hrs; 13 days). Consistent with human withdrawal expression, a subset of symptoms that included both somatic (e.g,. escape attempts, hyperactivity) and physiological (e.g., body temperature) items were still observed up to 2 weeks after the final heroin administration (see Figure 6G). In addition, the symptom expression was positively associated with the number of heroin infusions the animal took at the beginning of the study. Importantly, studies in which animals received forced opioid injections have reported hormone abnormalities (Li et al., 2010), persistent anxiety (Bravo et al., 2019), sleep disturbance (Khazan & Colasanti, 1972), and increased motivation to seek heroin (Bossert et al., 2015; Shen et al., 2011) during the protracted period. These behaviors were not fully examined here and would be important next steps to be explored using this paradigm.

Many of the symptoms evaluated throughout the acute and protracted phases also varied according to sex, including the only physiological rating collected here (body temperature). Phase 2 evaluation revealed that female rats evidenced significantly elevated body temperature across timepoints, prior to the onset of somatic signs (e.g., 35.09°C at 0 hours, compared to 36.91°C at 312 hours), whereas males initially showed lower body temperatures, followed by an eventual increase (e.g., 34.13°C at 0 hours, 33.96°C at 16 hours, and 35.98°C at 312 hours). These data suggest that for females, body temperature may be an important withdrawal sign during both the acute and protracted withdrawal period, though males may only show a signal on body temperature during the protracted period. These findings converge with other preclinical evidence that sex mediates some opioid effects (Huhn, Berry, & Dunn, 2018). Current human clinical data on this topic are limited to differences in treatment presentation between men and women (Huhn, Berry, & Dunn, 2019), neonatal abstinence syndrome following prenatal opioid exposure (Klaman et al., 2017), and menstrual cycle abnormalities and amenorrhea following chronic opioid exposure (Santen et al., 1975; Schmittner et al., 2005). Given evidence that ovarian hormones differentially impact nicotine-related withdrawal, craving, and relapse vulnerability in women (e.g., Carpenter et al., 2006; Franklin et al., 2008; Saladin et al., 2015; Smith et al., 2015), the prospective evaluation of differences in opioid withdrawal expression as a function of sex, as well efforts to further elucidate the role of sex and endogenously cycling hormones on opioid withdrawal severity, is warranted.

Strengths of these studies include the self-administration procedure, assessment of symptoms at baseline prior to heroin self-administration, and evaluation of outcomes in male and female rats. The study is limited by the lack of a placebo control group and reliance on a single animal strain (Long Evans rats), though there is no evidence that spontaneous withdrawal severity varies across rat strains (Cobuzzi & Riley, 2011). The access regimen used here was relatively short (3 hrs/day as opposed to 6 hrs/day) and inherently prevents animals from having uniform levels of opioid exposure, though it still produced quantifiable withdrawal symptomatology with variability that is likely similar to what is observed in humans. This study also summed symptoms according to the number of times they occurred, as opposed to human withdrawal assessment that often weighs the value of some infrequent symptoms more highly (e.g., vomiting is indicative of more severe withdrawal than nausea; Gellert & Holtzman, 1978). Weighing the symptoms according to their expected frequency would be a valuable next step for this line of research but was premature at this stage because the natural history of symptom incidence following these unique study conditions had not yet been determined. It is also not possible to determine exactly which preclinical symptoms correspond to human symptoms, though the symptoms did cluster into general domains that are conserved across species. Overall, these data provide an important foundation upon which future evaluation of withdrawals can work to refine the scoring system.

Additional studies that explore more features of this design are also warranted, including a direct comparison of precipitated and spontaneous withdrawal, evaluation of whether the context of drug self-administration alters withdrawal symptomatology (e.g., a saline substitution condition), comparison of outcomes following exposure to more than one unit dose or different patterns of opioid consumption (e.g., differences in duration of exposure, variability in day-to-day exposure patterns), and examination of outcomes in the context of hormone phase. Importantly, these data had notable outliers that may have impacted study results; these outliers were not removed from the analyses because there was no clear and justifiable reason to disregard them but additional studies that replicate these effects are needed. Finally, it will also be important that these questions be evaluated using other model (mice, nonhuman primates) species.

## Conclusions

The current results serve to provide a baseline for future research. Rats in this study were trained to self-administer heroin and then underwent a spontaneous withdrawal syndrome that was measured during an acute and protracted period. Outcomes showed variable incidence and duration of symptoms, many of which differed across males and females. Outcomes were also associated with rates of self-administration, which corresponds nicely with human opioid withdrawal. Since many of the symptoms that have been previously reported following forced opioid administration and naloxone-precipitated withdrawal in rats were observed at very low levels or not at all in this study, additional research that directly compares these two syndromes following heroin-self-administration during both the acute and protracted periods is needed. The vast majority of medications that show a signal for withdrawal suppression in animals do not translate into human studies (Dunn et al., 2019), and it is possible that medications that reduce the severity of a high-magnitude precipitated withdrawal may require higher doses or larger sample sizes to show an effect or register a signal for a less severe spontaneous withdrawal syndrome. These data expand the limited understanding of acute and protracted withdrawal syndromes in male and female animals following heroin self-administration, consistent with recent NIDA interest in enhancing existing animal models of SUDs to more closely approximate the human experience. Ultimately, these data provide a thorough and sensitive characterization of the natural history of an ecologically-representative model of human opioid behavior by focusing on spontaneous opioid withdrawal following heroin self-administration in male and female rats and support additional research to further develop this model.

## Acknowledgements

The authors would like to thank Robert Zaczek (NeurOp, Inc.), and Ngoc Van Do, Jose Piña and Vincent Carfagno for technical assistance. This work was supported by the National Institutes of Health Grant DA046266 (to Robert Zaczek), and DA046526, DA036569, DA036569-S1, DA044479, and DA045881 (to CDG), and the Arizona Alzheimer’s Consortium (to CDG).

